# Deep learning-based photo-identification for non-invasive monitoring of animal populations: Application to penguins and tortoises

**DOI:** 10.1101/2025.07.01.661353

**Authors:** Nicolas Durr, Miguel Rodriguez-Olmos, Julien Courtecuisse, Caroline Gilbert, Sylvie Laidebeure, Alexis Lécu, Ashley L Nord, Francesco Pedaci, Jason D Whittington, Yvon Le Maho, Víctor Planas-Bielsa

## Abstract

We describe DL-ID, a deep learning-based framework for the individual identification of animals from images with higher accuracy and characteristics that make it more robust in challenging conditions, such as field studies, than widely used algorithms. This is demonstrated through testing with small datasets involving two species that exhibit distinctive morphological features: Humboldt penguins (*Spheniscus humboldti* Meyen, 1834) and Hermann’s tortoises (*Testudo hermanni* Gmelin, 1789). We define a confidence index based on entropy within the model that significantly improves handling of the open set recognition problem, offering a new way to discriminate unknown individuals. DL-ID’s accuracy of 87%, even with as few as four images per class, contrasts with the usual practice in deep learning of building large datasets to assess performance. The model outperforms traditional photo-identification methods like Wild-ID and I3S Pattern, offering a significant advancement for research and conservation efforts. Its efficiency and adaptability indicate its potential in real-time monitoring, opening new possibilities for wildlife conservation. Instead of designing a complex new deep learning model, we focused on adapting existing methods to address a relevant ecological problem. Our approach effectively tackles issues like limited training data and recognizing new individuals in field studies. Notably, it performs well even with small datasets, making it particularly useful for data-limited ecological research. This makes our approach a valuable step forward in applying deep learning to ecological studies.

## Introduction

Individual identification is a crucial tool in life sciences and conservation for tracking population dynamics, assessing survival and reproductive success, and understanding behaviors and genetic patterns (Clutton-Brock & Sheldon 2010; Desai *et al.* 2022; Karczmarski *et al.* 2022; Schneider *et al.* 2019; Tuia *et al.* 2022). It is critical for collecting demographic data, such as survival rates, breeding success and first breeding age, and for facilitating capture-mark-recapture studies that inform conservation strategies and wildlife management policies (Ashe & Hammond 2022, Bolger *et al.* 2012; Clutton-Brock & Sheldon 2010; Gauthier-Clerc *et al.* 2004; Morrison *et al.* 2011).

Invasive techniques for individual identification, such as physical tags, pose ethical and practical concerns, inducing more stress from capture, potential harm and behavioral changes (Dugger *et al.* 2006; Gauthier-Clerc *et al.* 2004; Zemanova 2020). Such methods can also have long-lasting negative effects on animals, such as the reduction in breeding success and survival of penguins resulting from the increased drag at sea (Dugger *et al.* 2006; Gauthier-Clerc *et al.* 2004; Saraux *et al.* 2011). Capture for tagging can be particularly difficult and risky in endangered species (Bendik *et al.* 2013; Desai *et al.* 2022; Núñez-López *et al.* 2021; Zemanova 2020). RFID tags reduce some of these impacts by allowing subcutaneous implantation, thereby avoiding the impacts of external tags (Carter *et al.* 2014; Le Maho *et al.* 2014). However, they are still limited by very short reading distances, and they still require initial capture of the animals for tagging (Ferreira *et al.* 2020; Le Maho *et al.* 2014).

Photo-identification techniques rely on individual-specific skin patterns and eliminate the need for capture, offering a non-invasive alternative for wildlife researchers (Bauwens *et al.* 2018; Desai *et al.* 2022; Johansson *et al.* 2020). However, the increased photo collections present data processing challenges (Rigoudy *et al.* 2023; Tuia *et al.* 2022), including extended handling time (Schofield *et al.* 2019; Tuia *et al.* 2022), complex protocols (Choo *et al.* 2020; Elliser *et al.* 2022; Urian *et al.* 2015) and observer subjectivity (Ashe & Hammond 2022; Johansson *et al.* 2020; Urian *et al.* 2015). Software solutions like I3S Pattern and Wild-ID have been developed to streamline these processes and ensure data integrity (Bolger *et al.* 2012; Burghardt *et al.* 2004; Van Tienhoven *et al.* 2007). Yet, these systems require substantial researcher involvement (Givord-Coupeau & Rey 2023; Lorm *et al.* 2023) and strict adherence to specific photo criteria (Bendik *et al.* 2013; Matthé *et al.* 2017; Salom-Oliver *et al.* 2022). This functionally restricts the data variety (Ashe & Hammond 2022; Elliser *et al.* 2022; Urian *et al.* 2015) and impacts both the reliability and scope of studies (Choo *et al.* 2020; Elliser *et al.* 2022; Johansson *et al.* 2020; Karczmarski *et al.* 2022), particularly for the case of rare or social species aggregated in very large numbers (Ashe & Hammond 2022; Elliser *et al.* 2022; Johansson *et al.* 2020; Sherley *et al.* 2010).

Today, deep learning architectures, such as Convolutional Neural Networks (CNNs), mitigate many identification challenges, offering significant advancements in artificial neural network capabilities (LeCun *et al.* 2015; Vidal *et al.* 2021). Applied to animals, they outperform humans in various classification tasks (LeCun *et al.* 2015; Miao *et al.* 2019; Tuia *et al.* 2022), including individual recognition (Schofield *et al.* 2019; Wang & Deng 2021). CNNs excel particularly in image segmentation and recognition, identifying complex patterns with greater accuracy than humans (Clapham *et al.* 2020; Ferreira *et al.* 2020; Miao *et al.* 2019). They have a higher precision than traditional semi-automated methods (Bolger *et al.* 2012; Burghardt *et al.* 2004; Schneider *et al.* 2019; Van Tienhoven *et al.* 2007). However, directly comparing methodologies remains challenging due to inconsistencies in protocols (Karczmarski *et al.* 2022), photo processing (Urian *et al.* 2015; Vidal *et al.* 2021; Wang & Deng 2021), evaluation metrics (Liu *et al.* 2019; Vidal *et al.* 2021; Wang & Deng 2021), and reporting (Choo *et al.* 2020; Urian *et al.* 2015), and this is compounded by the scarcity of benchmark datasets (Ferreira *et al.* 2020; Liu *et al.* 2019; Vidal *et al.* 2021).

A notable limitation of conventional CNN models is their design as classifiers based on a fixed set of known individuals, which does not address difficulties in field applications that also require classifying novel individuals, leading to misclassifications (Ferreira *et al.* 2020; Gómez-Vargas *et al.* 2023; Vidal *et al.* 2021). This issue, known as the open set recognition problem (Safaei *et al.* 2024; Vidal *et al.* 2021), is critical for the reliability of population estimates (Ashe & Hammond 2022; Johansson *et al.* 2020; Morrison *et al.* 2011) and highlights a persistent challenge across both modern and traditional identification methods. This problem is compounded in traditional photo-identification studies, which may not consistently document or account for excluded pictures (Choo *et al.* 2020; Elliser *et al.* 2022; Johansson *et al.* 2020). Such omissions can introduce bias and uncertainty into the results (Ashe & Hammond 2022; Johansson *et al.* 2020; Urian *et al.* 2015).

An additional challenge inherent to CNNs is the required size of training datasets, which is usually large (LeCun *et al.* 2015; Liu *et al.* 2019; Wang & Deng 2021). In ecological contexts, acquiring a large number of images is generally impractical (Desai *et al.* 2022; Ferreira *et al.* 2020) — often comprising three or fewer samples per encounter — so it is imperative to meticulously examine how data scarcity impacts the efficiency and precision of the identification system (Gómez-Vargas *et al.* 2023).

Here, we introduce DL-ID, a new operational framework using CNNs designed to address the open-set recognition problem, while also accommodating the use of smaller datasets for the accurate and efficient recognition of individual animals. We demonstrate this framework application using the endangered Humboldt penguin, a species with distinctive chest spots for which there is an urgent need for more biological knowledge and a reliable evaluation of its conservation status (De la Puente *et al.* 2013; Mcgill *et al.* 2022). We further demonstrate the adaptability of our approach by applying it to Hermann’s tortoises, another species with more subtle yet individually unique features. The successful application of our method to these diverse taxa highlights its versatility and underscores its potential to contribute to wildlife conservation and ecological research across a broad spectrum of species. Instead of designing a complex new deep learning model, we focused on adapting existing methods to address a relevant ecological problem (Appendix 1). Our approach effectively tackles issues like limited training data and recognizing new individuals in field studies. Notably, it performs well even with small datasets, making it particularly useful for data-limited ecological research. This makes our approach a valuable step forward in applying deep learning to ecological studies.

## Material and methods

### Humboldt Penguins

#### Animals and data collection

The study took place in France at the Paris Zoological Park (hereafter PZP, 48° 50’ N, 2° 24’ E). At the project beginning in February 2021, the Humboldt penguin colony was composed of 24 individuals: 7 females and 17 males. At its end in June 2023, it had grown to 31 individuals, corresponding to 14 females and 17 males. Data acquisition was conducted from outside the enclosure during three sessions of 5 months, covering the breeding season. Besides incubating and breeding at nest, penguins are active both spatially and temporally. Photo sessions were randomly performed at different dates between 9:00 am and 4:30 pm, instead of following a repeated schedule. Those elements were important to avoid introducing biases in datasets, as the PZP penguins may have adopted some routines, and it permitted data acquisition across a greater range of environmental conditions (Figure 1A-B).

**Figure 1.**
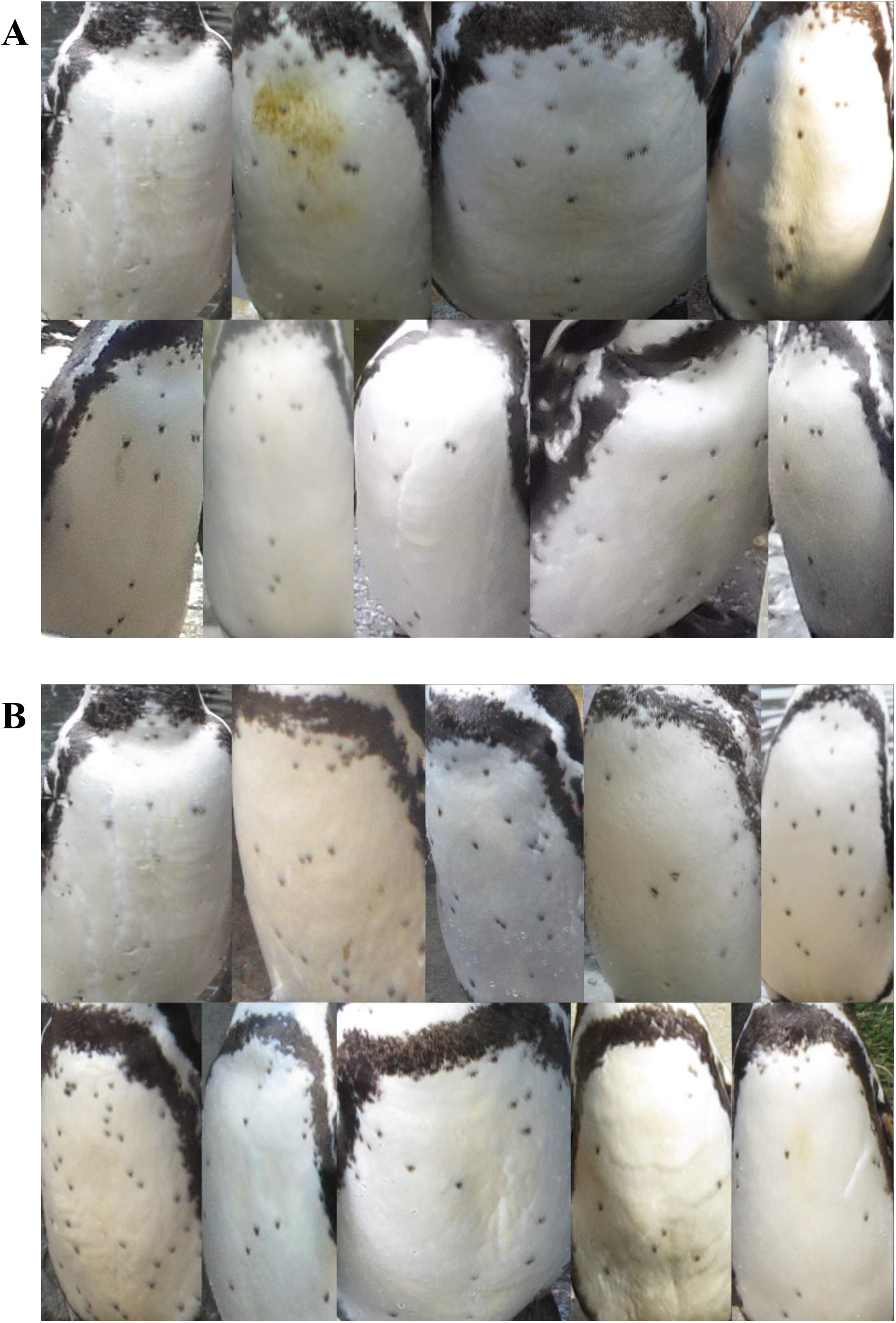
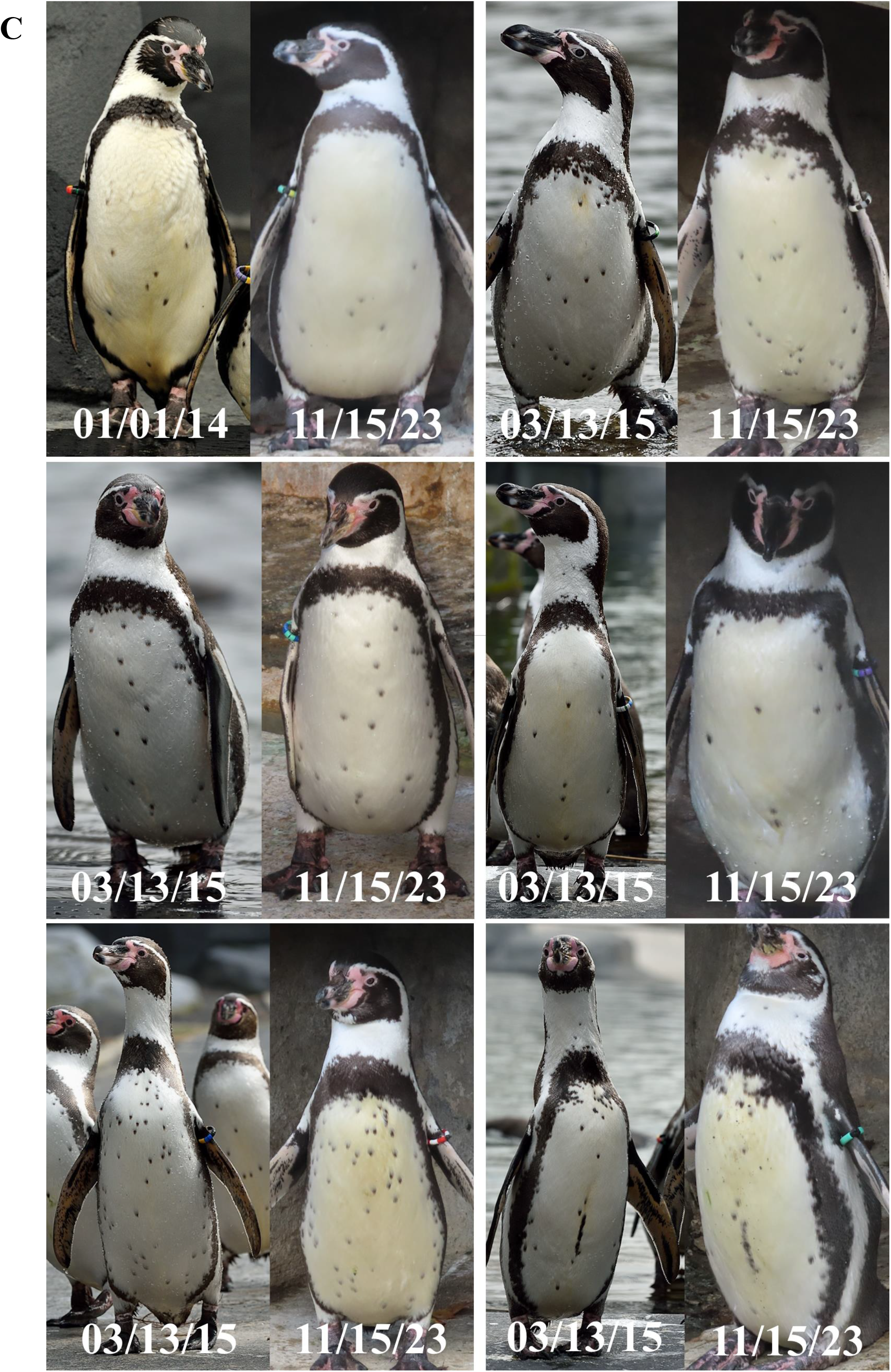
Overview of the data collection features. Intra-individual variability (A). Environmental heterogeneity introduced in the sampling. Inter-individual variability (B). Diversity of spot patterns and their distinctiveness. Pattern stability (C). Evolution for six penguins between their oldest record (©MNHN-François-Gilles Grandin) and their most recent picture.

#### Persistence of the spot pattern

As a mandatory step for photo-identification investigations, we assessed that the chest pattern complies with a strict requirement of stability. Our team was given access to the Museumedia library (https://www.mnhn.fr/en/museumedia-the-multimedia-collection), gathering media files from the National Museum of Natural History (MNHN). The oldest photographs from the Humboldt penguin colony at the PZP were taken in 2014 (©MNHN -François-Gilles Grandin), and they attest that no significant physical changes occurred over a decade (Figure 1C).

#### Construction of datasets

To have an objective comparison between our deep learning approach and existing methods, i.e. SIFT and SURF (Liu *et al.* 2019), we created a standard benchmark database with images of 26 Humboldt penguins, with known identities based on the uniqueness of their chest patterns (Burghardt *et al.* 2004) and color rings. Picture classification per individual was double-checked by two independent observers to ensure the database was free of labeling errors. Penguin chests were cropped by hand to remove most background (Johansson *et al.* 2020; Lorm *et al.* 2023) and unmarked body parts (Bolger *et al.* 2012), and special care was made to remove the color band to avoid potential bias during the analyses (Ferreira *et al.* 2020). All images were resized to 200 × 300 pixels.

The database comprised between 42 and 58 images per penguin. It was subdivided into three non-overlapping groups: a test set containing 20% of the images per individual, a validation set of the same size and a master training dataset denoted P_60%_. For some experiments, this master training dataset was reduced to smaller datasets, as further explained in the following sections. We also used the dataset P_80%_ which corresponds to the final training pass, i.e. P_60%_ dataset plus validation set. The validation dataset was utilized to monitor the model performance with respect to hyperparameters. The test set was reserved for a single use after the model was finalized, and the results from this test set are those reported. Before their use, all the images were packed into a numeric NumPy array of size 1630 × 200 × 300 × 3px where the last dimension corresponds to the color image having three channels. A metadata file contained all the image names, labels and belonging datasets.

### DL-ID construction and tuning

#### Architecture and initialization

The main approach to obtain a convolutional architecture for our model consisted in trying designs whose performance was acknowledged on the ImageNet challenge (https://www.image-net.org/challenges/LSVRC/index.php). These networks have been designed to succeed in very general-purpose image recognition problems, and can deal with almost any type of image, since they are usually trained with the ImageNet database, a representative sample of the visual diversity of the world. Typically, and mostly for computing reasons, these networks are used with the parameters obtained from the ImageNet training, and then the top classification layer is substituted with a custom layer whose parameters are trained ad hoc for the new classification problem. This is known as transfer learning (Ferreira *et al.* 2020; Schneider *et al.* 2019). A variation on this theme is fine-tuning, which consists in retraining some of the last layers before the top one. In both approaches, most of the deepest layers remain unchanged with the same parameters obtained during the ImageNet training.

In our approach, we have used the network architectures, and have fully trained them from random initialization. This is computationally expensive at the training stage, but not at the inference stage, i.e. validation and testing steps. With modern GPUs, a full training of most tested networks is doable. In our case, we have used the available GPUs in Google Colab (https://colab.research.google.com). A major advantage of this strategy is that every parameter of each layer tailor the individual penguin recognition problem. Since most of the deeper layers of a CNN adapt themselves to recognize the most abstract and basic features of the images, a very general-purpose model proven to successfully classify virtually any image in the world into the right category will be a good candidate for our goal.

Several architectures were utilized, namely InceptionV3, InceptionResNetV2, VGG19, MobileNetV2, ResNet152V2, ResNet50, NASNetLarge and MobileNet. For each architecture, we conducted experiments with grayscale/color images as well as different image and batch sizes. We found no need to tune the learning rate, likely due to our use of the Adamax optimization algorithm, which automatically adjusts the learning rate during training, with an initial value of 0.001 by default. Extensive experimentation was also carried out regarding the number of epochs, and during the validation analysis we settled on 500 epochs except for the P_2_ experiment, where 800 epochs were needed due to the extremely low number of images available. Overfitting was not observed in any case, and the training reached a plateau in every iteration. Ultimately, we selected the InceptionV3 architecture with a batch size ranging from 4 to 16 depending on the particular problem and three-channel color images, maintaining the original size of 200 × 300px. Additionally, we experimented with different levels of image augmentation, including rotation, horizontal and vertical shifts, brightness adjustment and contrast enhancement, a general procedure that helps to design models robust to ecological scenarios (Ferreira *et al.* 2020; Liu *et al.* 2019), especially in a few-shot context (Gómez-Vargas *et al.* 2023).

#### Implementation of ensemble method

For inference, we used an ensemble approach which consists in combining the results of models trained independently (Malik *et al.* 2021; Mohammed & Kora 2023). We trained five instances of the selected foundational network (InceptionV3) with a different random initialization each time. This way, each model reaches a different local minimum of the categorical cross-entropy loss function. The prediction of each model is a probability vector, with its dimension representing the number of known classes in each species. For each individual prediction, the sum of the components of its probability vector is 1. By averaging for every penguin over its five corresponding probability vectors, we get a new probability vector that does not come from any of the networks in particular but from their aggregation. Finally, the predicted class of the individual was determined using the argmax function on the ensemble probability vector. Alternative aggregation methods exist, such as majority vote by letting each instance predict a category and then looking for the most predicted category (Mohammed & Kora 2023). However, we found that the accuracy of the average ensemble was higher in all cases.

#### Confidence Index for open set problem

In another experiment, we introduced unknown individuals at the inference stage to test if our ensemble model could detect an extraneous animal that was unknown during training, and therefore did not have a corresponding category in the output probability vector. Obviously, applying the ensemble model to an image of this unknown individual will still produce a probability vector that, after the argmax probability collapse, would incorrectly identify this as a known individual. Our goal was to exclude this possibility and have a mechanism that flags this output vector as corresponding to an unknown individual.

To achieve this goal, we defined a Confidence Index (CI) as one plus the normalized entropy of the predicted probability vector, that is:

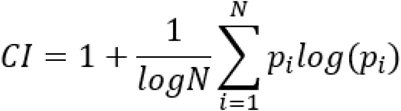

where p_i_ represents the probability value assigned to each class by the ensemble model, and N is the total number of classes in which each instance of the foundational model has been trained. It is important to note that when p_i_=0, the term p_i_ log(p_i_) is defined to be 0. This index gives a value of 1 if there is class for which the probability is 1 (meaning that the remaining components of the ensemble probability vector are 0) and 0 when each class has equal probability, i.e. p_i_=1/N, and therefore the uncertainty is maximal. The combined use of the ensemble model and the CI for flagging unknown individuals is explained below.

### DL-ID comparison with other methods

The performance of our deep learning approach was compared to I3S Pattern (https://reijns.com/i3s/i3s-pattern/) and Wild-ID (Bolger *et al.* 2012) (see details in Appendix 2). Two different experiments were designed to evaluate their identification capabilities: recognition of known individuals (***Known***), which includes the resilience of the methods to the decreasing size of the training dataset, and discrimination of unknown individuals (***New***). For each experiment, the training and validation datasets can be used to optimize the model, but the results reported for the test dataset can only be used at the end of the process, when no further hyperparameter tuning or modification of any kind will be applied to the model. Following standard practice in deep learning, the training set was used to train models and learn their parameters, while the validation served as a guide to fine-tune hyperparameters and decide on the number of epochs. In a similar way, I3S Pattern metadata were optimized by comparing the validation set against the training database (Appendices 3 and 4), but the user interface of Wild-ID does not allow any modification of internal parameters.

#### Recognition of known individuals

We designed the ***Known***experiment to evaluate the accuracy of the recognition methods for known individuals under different training sizes. For this, we used a series of nested training datasets, each with fewer images than the previous one, using images of 19 penguins. To create the best possible deep learning model, we first used the entire training dataset gathering 6 to 35 images depending on the individual in an experiment named **P_60%_**. Seven subsequent experiments denoted **P_50%,_ P_40%,_ P_30%,_ P_20%,_ P_10%_ P_5%_**and **P_2_**were conducted using less training data, but were always evaluated on the same validation and test datasets. Each training dataset was constructed in a nested way as a random sub-selection of the previous one, and its name indicates its size. For instance, **P_60%_**contains 60% of the dataset as training material, and **P_10%_**contains 10% of the images. The smallest experiment called **P_2_**was trained using only two images per individual.

In typical deep learning practice, one would train on the training set, use the validation set for tuning hyperparameters and/or decide among various models. Before reporting the final results, the model is retrained using all the available data, which includes both the training and validation set. We could not do this since that would interfere with checking the resilience of the model with respect to the size of the training set. However, we define an additional experiment, denoted as **P_80%_**, as follows: it was conducted using the best configuration determined from the preceding stage **P_60%_**after validation. In fact, we merged the validation dataset, representing 20% of the images, with the **P_60%_**training database for a comprehensive final training iteration.

Assuming the absence of unknown individuals and labeling errors, we evaluated the performance of identification methods according to two metrics: total accuracy for a dataset and time spent per image prediction at inference. For the evaluation of accuracy, only top-rank accuracy was considered. This means that accuracy was measured based on the model’s most likely prediction. This metric was selected as it is a standard metric used to evaluate the performance of deep learning models (Schneider *et al.* 2019; Wang & Deng 2021). This approach contrasts with some ecological studies, which consider a match positive if the individual is among the ten most likely predictions (Givord-Coupeau & Rey 2023; Lorm *et al.* 2023), but also many others referring to even higher matching ranks, i.e. 20, 30 or 100 (Bendik *et al.* 2013; Matthé *et al.* 2017; Suriyamongkol & Mali 2018). In practice, analyzing many potential matches requires repetitive manual comparisons, a tedious and error-prone approach (Schneider *et al.* 2019; Tuia *et al.* 2022) already attested as highly variable (Johansson *et al.* 2020; Horn *et al.* 2014) and likely associated with underestimated error rates (Johansson *et al.* 2020; Urian *et al.* 2015). Due to the lack of availability of large training sets, cross validation was not performed and is not included in the results reported.

Concerning processing time, we did not include image pre-processing, as labeling, cropping and random repartition steps were common to all tested recognition methods. We also did not include in this analysis the ‘fixed cost’ of setting up the hyperparameters correctly for I3S Pattern and our model, as this is done only once for the species considered. The time at inference was assessed from the data input to the generation of output files. Note that this metric was calculated for a pilot study whose subject identities were known, so identifications were validated automatically by relying on picture labels to gain significant time.

#### Discrimination of unknown individuals

For the testing with ***New***individuals, the procedure remained the same as for the ***Known***experiment, except for the incorporation of new identities and the change of evaluation metric. The training dataset was the one used for the **P_80%_**experiment, but 171 additional images of seven new penguins were inserted in the validation test before optimizing DL-ID and I3S Pattern parameters. Then, in addition to the standard test dataset, we included the same number of images for those seven individuals that the network had never seen before. The identity displayed on those new pictures was double-checked beforehand using both individual chest and band, so they did not belong to any penguin in the training dataset, and therefore did not have allocated categories.

As already explained, the inference method outlined before, in which the network produces a probability vector of the same dimension as the number of known individuals and the argmax function is applied afterwards, will predict that this new individual is one of those present in the training set. However, if the individual is new and our ensemble method is accurate the ensemble probability vector will concentrate around a vector with a 1 in the position corresponding to the ground truth of the individual and 0 in the rest. The previously introduced CI, which is normalized between 0 and 1 is effectively a measure of the entropy of this probability vector and has 0 in extreme cases (when each position has the same value 1/N, or every possibility gets the same probability) or 1, in the ideal case when the model predicts a class with probability 1, since then the rest of the entries of the probability vector must be 0. In our problem, we set a threshold for CI and declare that those probability vectors with CI less than this threshold correspond to new individuals, and ignore the inference obtained after the argmax step. We point out that the use of entropy, or some variations, like the one we propose here, is a very natural idea to handle the open set problem (Safaei *et al.* 2024). Since we are more interested in determining whether an individual is known or new, rather than recognizing specific individuals, we slightly adjust the power of the combined use of multicategory training with the CI. This adjustment allows us to frame the task as a binary classification problem, where 0 corresponds to a known individual regardless of their identity and 1 corresponds to a new individual.

To test the quality of the discrimination between new and known individuals, we use the Receiving Operating Characteristic (ROC) curve instead of binary accuracy, as it represents a classical tool in machine learning to evaluate binary discrimination tasks simultaneously for all possible thresholds (Sherley *et al.* 2010; Wang & Deng 2021). In our DL-ID framework, normalized entropy (CI) was considered as a potentially relevant discriminator, whereas, for I3S Pattern and Wild-ID, the similarity score computed for the top match acted as such. For each threshold value, the true positive and false positive rates were calculated, corresponding respectively to the proportion of pictures correctly and wrongly attributed to a new identity. The ROC curve compiles all those results on a single chart, allowing to visually appreciate the discrimination efficiency. Then, the area under the ROC curve (AUC score) can be computed to quantitatively assess model performance. The AUC score lies between 0 and 1; a value of 0.5 corresponds to a completely random prediction of the outcome, while a value of 1 corresponds to a perfect prediction. This approach encompasses all the possibilities of computing binary accuracy for each possible choice of CI threshold. In particular applications and depending on our interest on controlling the number of false positives or negatives, we could use this for setting an optimal CI threshold, although this specific issue has not been treated in this work and is case-dependant.

### DL-ID transferability to Hermann’s Tortoises

#### Animals and data collection

This part of the study took place in France at the “Station d’Observation et de Protection des Tortues et de leurs Milieux” (SOPTOM, 43° 18’ N, 6° 12’ E), where captive Hermann’s tortoises are located. We took images of 24 individuals. Since the individuals were not marked, we followed a specific procedure to ensure accurate identification. We captured each individual and placed it on a brown-colored cardboard square in a separate location away from the main area. The same cardboard was used for all individuals to avoid background bias. All images of each individual were taken in a single session, and all individuals were photographed on the same day under consistent light and climate conditions. This ensured uniformity across the dataset.

We collected dorsal images of each tortoise shell in at least two different settings, i.e. shadow and sun, and used a variety of focal lengths to capture diverse images. This approach mimicks the high heterogeneity from field conditions and contrasts with previous studies (Carpentier *et al.* 2016; Carter *et al.* 2014; Salom-Oliver *et al.* 2022; Suriyamongkol & Mali 2018). Approximately 100 images were taken of each individual. For each one, we created a separate folder in the camera, assigning arbitrary distinct names to ensure that images from the same individual were collected in the same folder. For each folder, we added an arbitrary and distinct label name, and all images for the same individual contained the label as a prefix and a sequential number to differentiate the images uniquely. This allowed us to label the dataset with absolute certainty, as all individuals could be accurately identified in the folder hierarchy.

#### Construction of datasets

After inspection, we removed all images in which a significant part of the individual was outside of the frame, as they moved during the photo collection. After the incomplete images were discarded, we applied a simple background removal algorithm in python and rescaled the images to 300 × 300px. For some images, the background removal was not perfect, and some shadows were retained, but we left these images in the active database to demonstrate that precise and time-consuming image preprocessing is not needed (Appendix 5). Finally, to make a fairer comparison between Humboldt penguin and Herman tortoise, we randomly selected 50 images of each individual of which 30 were allocated to the training set, 10 to the validation set and another 10 to the test set. This setup is equivalent to the known P_80%_ experiment for the Humboldt penguins.

#### Recognition of known individuals

We followed a similar procedure as for the Humboldt penguins. Using color images, we first explored several architectures from our list of foundational models. For the Hermann’s tortoise, we used again the InceptionV3 architecture. We then explored different batch sizes, retaining the model with a batch size of 16, as it exhibited less variation. We did not explore different image sizes nor color/greyscale, because the resolution is clearly enough to show the shell patterns, and the used GPUs could handle all the images of our training dataset. All models were trained for 1000 epochs, and, as with the Humboldt penguin, overfitting was not observed in any case. Once the ‘best’ model was selected, we created four additional initializations, resulting in five instances of the model for our ensemble predictor. We used the average probability for aggregation and applied the argmax function to this probability vector to infer the most likely class.

## Results

### Classification of known and unknown identities

The new framework, DL-ID, is a deep ensemble model, composed of multiple parallel instances of identical CNNs (Figure 2). Each of these instances is based on the well-known InceptionV3 architecture. Until recently, it was one of the most successful architectures in the universal image recognition challenge ImageNet (https://paperswithcode.com/sota/image-classification-on-imagenet). Several alternatives with good results in ImageNet have been tested (see methods) and InceptionV3 was selected as a compromise between its complexity, i.e. training efficiency, and performance for our problem. DL-ID provides two outputs: first, it determines the most likely match for the individual in the image from a pool of known individuals. Secondly, and crucially, it generates a metric termed the Confidence Index (CI), which assesses the likelihood that the individual is already in the training dataset or corresponds to an unknown entity (Figure 2).

**Figure 2.**
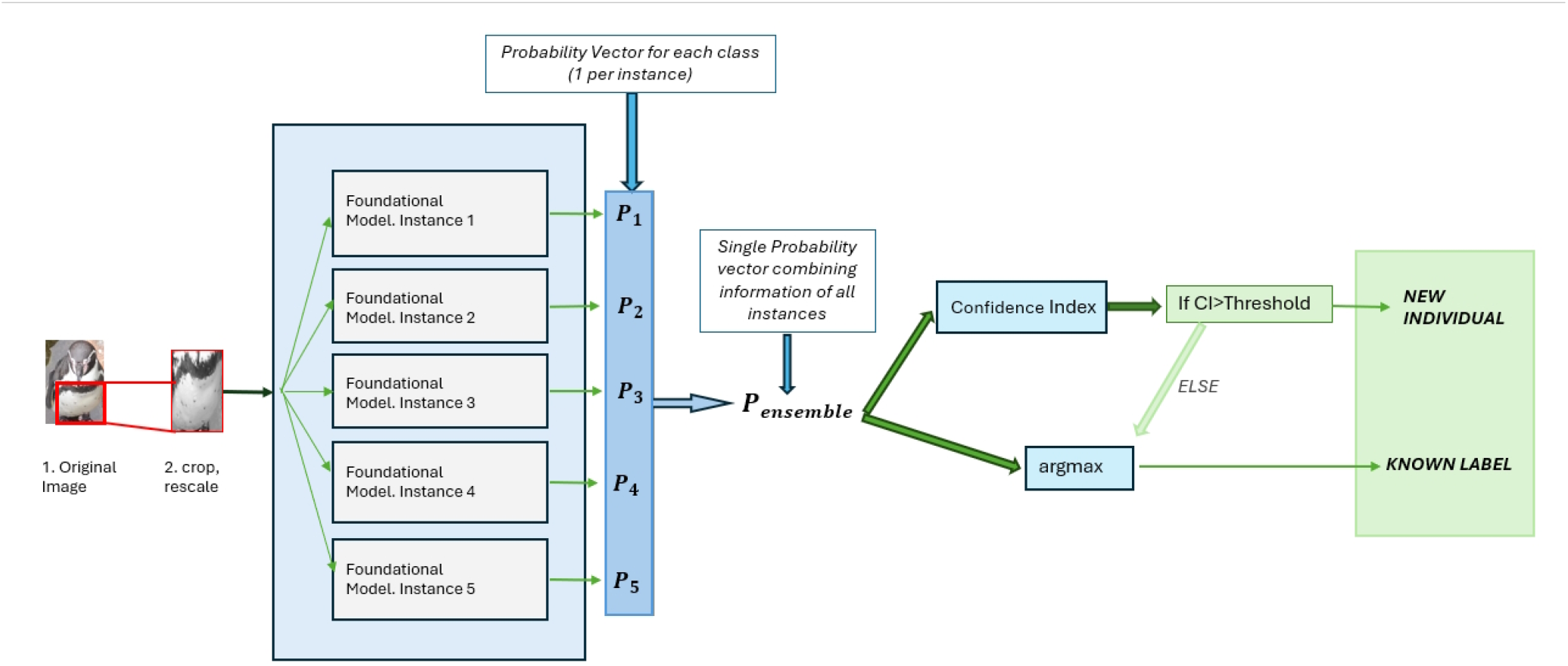
DL-ID architecture and processing steps. The ensemble model combines five models. Each has the same architecture but a different initialization and is trained independently on the same dataset. During inference, each image, after cropping and rescaling, is analyzed by all models, generating a probability vector for potential labels. These probability vectors are averaged to determine the ensemble probability for each label. The most likely class is then identified using the standard argmax function. The CI is calculated from the average probability vector, and a new identity is predicted if it exceeds a pre-defined threshold. Otherwise, the picture is attributed to the individual selected by the argmax function.

The model was tested using varying proportions of our dataset as the training set. These training scenarios are labeled according to the amount of data used for training. We also provide a comparative analysis between DL-ID and two pre-existing methods implemented in Wild-ID and I3S Pattern (see methods).

### Humboldt penguins’ identification

A prerequisite for accurately identifying individual animals lies in the long-term stability of their distinguishing marks, regardless of external factors such as annual molting in birds. Our photo database of Humboldt penguins of known age, spanning three breeding seasons, includes individuals that underwent two complete plumage renewals. Despite these changes, our DL-ID model effectively identified individuals in various conditions, such as different angles, postures, distances, and times of day (Figure 1A). This demonstrates the model’s ability to recognize the consistent and highly distinctive visual features of individual penguins (Figure 1B). We also observed no significant changes related to the chest spots of six Humboldt penguins from images archived over a decade, indicating that training data from one year can be effectively used for identification in subsequent years (Figure 1C).

### Framework transferability to Hermann’s tortoises

After trying several model architectures, InceptionV3 was also selected as the most effective for Hermann’s tortoises, providing ideal conditions for a direct comparison with the identification rate obtained for Humboldt penguins. Then, specific training and validation were performed to fine-tuned models to recognize tortoise shell features, less distinctive compared to Humboldt chest spots (Appendix 5).

To be consistent with the Humboldt penguin experiment, Hermann’s tortoise datasets are built using the same data proportions, i.e. training, validation and testing datasets respectively at 60%, 20% and 20%. After tuning the hyperparameters, the training and validation datasets were merged for a final training pass. When trained on our largest dataset combining those two sets (P_80%_ = 763), DL-ID achieves an accuracy of 99% on the test set of Humboldt penguins (Table 1). In Hermann’s tortoises, we obtained an accuracy of 99.7%, using overhead pictures of their shells without capturing them.

**Table 1.** Training datasets and final accuracy. Balance between identities is optimized in small training sets to prevent overfitting. Identification accuracy is given for Wild-ID, I3S Pattern for direct or optimal comparison (Appendix 7) and DL-ID with or without data augmentation.

| Dataset name | Training images per individual |  | Previous methods |  |  | Our methods |  |
| --- | --- | --- | --- | --- | --- | --- | --- |
|  | min | max | Wild-ID | I3S Pattern direct | I3S Pattern optimal | DL-ID no aug | DL-ID aug |
| P<sub>2</sub> | 2 | 2 | 0.36 | 0.49 | 0.4 | 0.63 | 0.62 |
| P<sub>5%</sub> | 3 | 4 | 0.4 | 0.53 | 0.79 | 0.79 | 0.87 |
| P<sub>10%</sub> | 5 | 6 | 0.41 | 0.59 | 0.86 | 0.90 | 0.93 |
| P<sub>20%</sub> | 9 | 12 | 0.45 | 0.66 | 0.88 | 0.97 | 0.98 |
| P<sub>30%</sub> | 13 | 18 | 0.53 | 0.79 | 0.89 | 0.97 | 0.99 |
| P<sub>40%</sub> | 17 | 24 | 0.56 | 0.79 | 0.9 | 0.97 | 0.99 |
| P<sub>50%</sub> | 21 | 29 | 0.56 | 0.8 | 0.93 | 0.98 | 0.99 |
| P<sub>60%</sub> | 26 | 35 | 0.58 | 0.8 | 0.93 | 0.98 | 0.99 |

### Comparison with Wild-ID and I3S Pattern

In our study, with its accuracy of 99% for our Humboldt penguin benchmark dataset (P_60%_), DL-ID consistently outperforms previous photo-identification methods across varying training set sizes (Figure 3A). Based on the same images, Wild-ID achieves an accuracy of 58% and I3S Pattern reaches 80%. When following user guidelines and applying a precise chest cropping, I3S Pattern accuracy improves to 93%. However, this improvement comes at the cost of a significant time investment required for carefully selecting the region of interest on each image (Appendices 6 and 7). More accurate, I3S Pattern optimal setting was selected for performance comparison with DL-ID and Wild-ID in following experiments.

**Figure 3.**
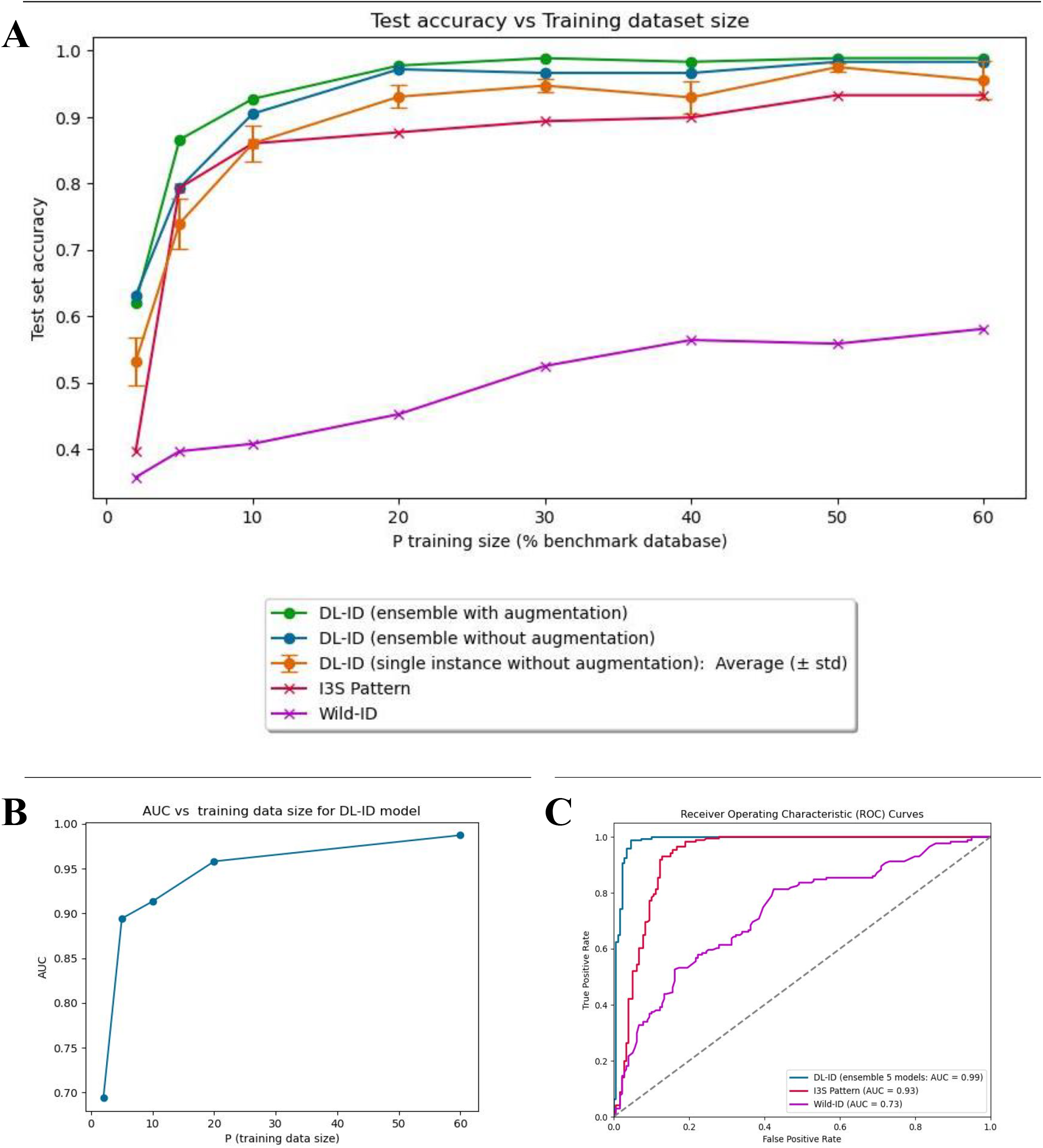
Performance comparison between methods. Identification accuracy of known individuals for different sizes of the training dataset (A). Implementation of data augmentation and ensemble approach increase identification accuracy, particularly with limited training material. Impact of training size on DL-ID ability to distinguish new individuals (B). ROC curves referring to the discrimination between known and new individuals (C). Compared to DL-ID top-notch performance, I3S Pattern and Wild-ID display lesser categorization capabilities, even if I3S Pattern achievement is already excellent.

For smaller training datasets (Table 1), DL-ID maintains high accuracy until P_10%_, corresponding to 10% of the database (103 images out of 942), after which accuracy begins to decrease. At P_5%_, it still achieves 87% accuracy, and at P_2_, i.e. with exactly two pictures per penguin, accuracy drops to 62%. In contrast, Wild-ID and I3S Pattern consistently show lower accuracies than DL-ID, and, as the training dataset decreases, they experience a steeper drop in performance (Figure 3A).

### Open set problem and unknown individuals

For each individual classified by DL-ID, a CI is calculated (see methods). The CI is used to decide whether an image corresponding to a known individual represents a new individual, i.e. not present within the training dataset. The Receiver Operator Characteristic (ROC) curve at figure 3B shows that DL-ID performance in the binary classification of individuals, as either new or known ones, increases rapidly with the database size. Relying on its CI to act as a threshold parameter, DL-ID handles the task better than the two traditional methods, as confirmed by figure 3C displaying the ROC and the Area Under the Curve (AUC) for different sizes of training dataset. Indeed, DL-ID efficiency only drops to the level of I3S Pattern after reducing the training size to 5%, representing 3-4 training pictures per penguin (Figure 3B).

Figure 4A compares the distribution of DL-ID CI values and similarity scores computed by Wild-ID and I3S Pattern. This visualization clearly demonstrates the effectiveness of the ensemble procedure, i.e. model averaging approach, in separating these distributions, thereby increasing the model ability to accurately distinguish between known and unknown individuals. DL-ID exhibits exceptional performance in detecting positive cases, that is identifying unknown individuals in the context of photo-identification. This high sensitivity, also referred to as recall, is a valuable feature in our methodology. In practical terms, prioritizing the capture of all positives, even if it results in a few false positives, ensures that no potential new individuals are missed during the identification process (Figure 4B). This allows to only include individuals with certain recapture history in analyses and offers a second chance to check a much smaller set of images, searching for a few remaining matches.

**Figure 4.**
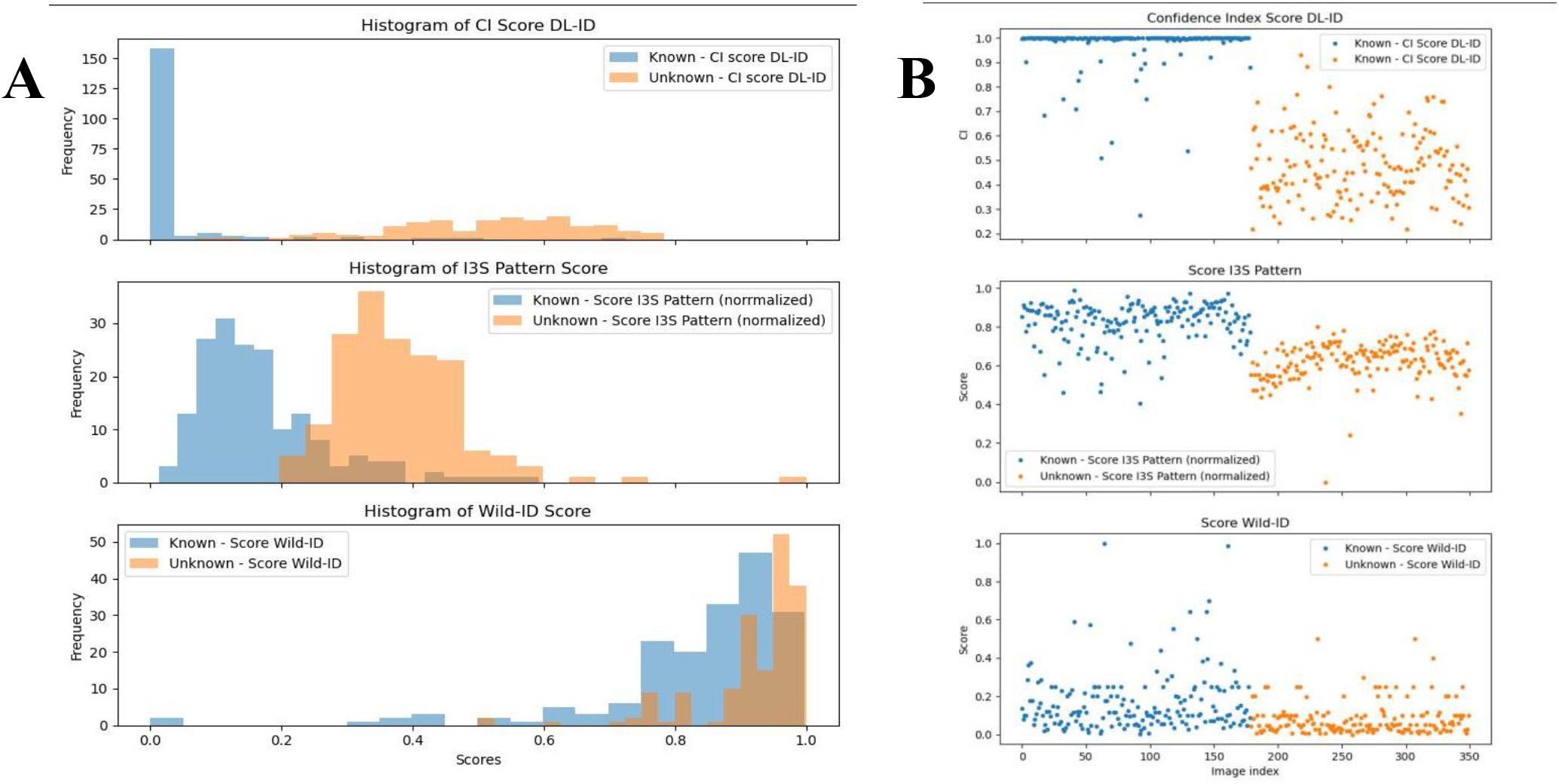
Discriminatory power of CI and scores. The test set combines the one from the Known experiment with another of equal size, consisting of unknown identities. Histograms of CI values and scores used to discriminate unknown identities (A). With its CI, DL-ID clearly separates known and unknown individuals. Another representation of the CI efficiency (B). Each point corresponds to a given image, its index being given on the x-axis. The first 179 images (blue) correspond to known individuals, while the second half (orange) corresponds to the unknown individuals.

### Time to process batch of images

The three photo-identification approaches also differ in terms of handling time and processing duration related to the images in the training database and the 179 testing pictures (Figure 5A). We differentiate here between the ‘training’ preparation phase, only performed once, and the inference phase, which is linear with the number of images to process.

**Figure 5.**
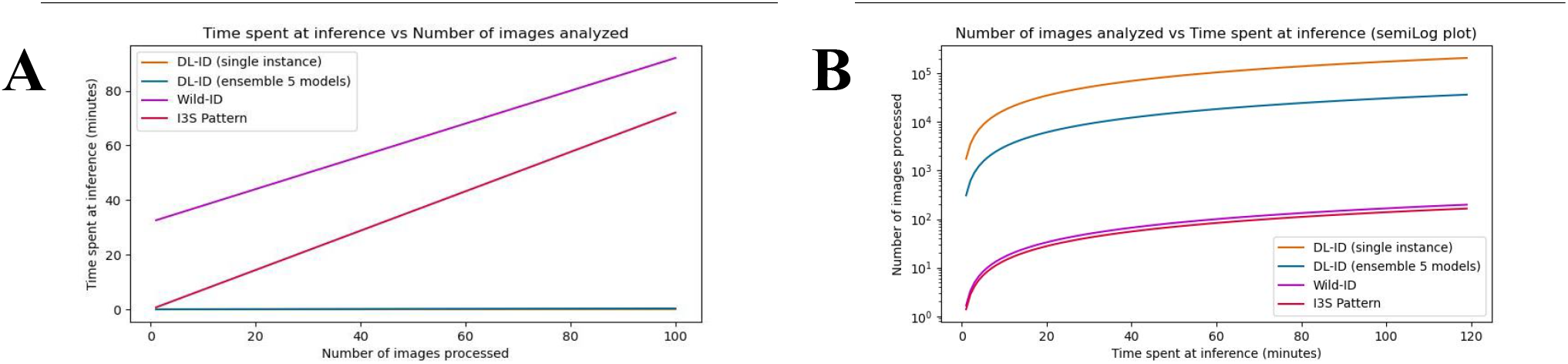
Performance in processing time. Time spent at inference per image analyzed (A). Wild-ID and I3S Pattern require time-consuming manual steps, posing a problem of scalability for large datasets. The single and ensemble instances of DL-ID appear of identical speed by superimposition, but single instance is faster. Number of images processed for a given time (B). In one hour, DL-ID can process more than 15,000 new pictures, whereas only 46 are processed with Wild-ID and 83 using I3S Pattern. While slightly less accurate for small training sets, a single model instance performs faster than the ensemble model, making it more appropriate for embedded systems.

For Wild-ID, the automated batch comparison of testing set against the training database take between 45 and 55 minutes. After feature extraction, user selection of the correct match, when displayed, typically takes less than 30 seconds per image, averaging 12 seconds when individuals are familiar. Therefore, after taking around 10 minutes to manually go through all training pictures, the review and matching of the remaining testing pictures take approximately 36 minutes for each identification experiment.

For I3S Pattern, three reference points and the region of interest for each image in the training database must be selected by hand (Appendix6), requiring between 40 and 60 seconds per picture and 43 on average. Thus, the total time required for training is about 8.5 hours. At inference, reference points must also be added, which requires about two hours for a testing set. In addition, there is a non-negligible amount of time spent for the batch comparison step depending on the metadata settings. While we are using an optimal setting of 18 points, I3S Pattern takes approximately 1 minute to compare between all paired pictures; the same procedure could last up to 30 minutes if 34 points are required as for the direct comparison experiment (see methods).

Our DL-ID model was trained on a T4 GPU, and each instance was trained in about 20 minutes, totalling close to 1.5 hours for the five instances in the ensemble. At inference time, once the model has been loaded into memory, it takes approximately 10 seconds to process the 179 images of the testing set on a standard laptop without GPU (Figure 5B).

## Discussion

DL-ID is a deep learning model based on CNNs that produces faster and more accurate identification of individual animals from photos than other tools currently being used in animal studies. Efficient training with minimal data and achieving open-set recognition remain two major challenges that continue to defy most investigations and even the most powerful CNNs. Our new framework addresses these challenges by offering an improvement over widely used tools, expanding its potential application in the field. Additionally, its robustness against environmental heterogeneity and its transferability across highly distinct features and species represent valuable contributions towards this objective.

In addition to the challenges directly examined, our study confirms the stability of the Humboldt penguin chest pattern over a span of at least ten years, as inferred from opportunistic observations of African penguins (Burghardt *et al.* 2004). While avian species may undergo gradual and subtle changes in their plumage features through successive molts during adulthood (Ferreira *et al.* 2020), this hypothesis has not been explored by the previous photo-identification studies focused on birds (Burghardt *et al.* 2004; Ferreira *et al.* 2020; Núñez-López *et al.* 2021). Evidence from other species suggests that aging can affect the reliability of identification (Horn *et al.* 2014; Schofield *et al.* 2019; Suriyamongkol & Mali 2018), and various factors such as melanophore size (Bendik *et al.* 2013), internal and abiotic conditions (Matthé *et al.* 2017), injuries and recovery (Ashe & Hammond 2022; Elliser *et al.* 2022) impact visual patterns at different rates. These findings highlight the necessity for a thorough investigation into how these factors influence long-term identification accuracy, regardless of the recognition methodology employed (Bolger *et al.* 2012; Carpentier *et al.* 2016).

A full technical comparison of our framework with the literature is difficult due to variations in protocol designs and discrepancies among studies. Notably, the number of potential matches manually analyzed can differ radically (Lorm *et al.* 2023; Meenakshisundaram *et al.* 2021; Salom-Oliver *et al.* 2022; Suriyamongkol & Mali 2018), affecting the overall accuracy reported, species-specific reliability and adaptability of the different software programs. Yet, our experiment, designed to remove these variables as factors, demonstrates that DL-ID surpasses its well-established counterparts, Wild-ID and I3S Pattern, in accuracy by an additional 35% and 18% on our benchmark dataset (P_60%_). Furthermore, DL-ID exhibits significantly improved time efficiency, processing 200 images 190 and 178-fold faster, respectively. This superior performance is maintained even with smaller training datasets, demonstrating DL-ID robustness.

Our demonstration of the efficacy of DL-ID in the identification of individual Humboldt penguins, including both known and unknown identities, illustrates its potential utility in ecological and conservation studies (Ferreira *et al.* 2020; Gómez-Vargas *et al.* 2023) to monitor wildlife with a lower impact than most current field campaigns. Indeed, avoiding specimen captures should be a paramount consideration from ethical, scientific and conservation perspectives (Desai *et al.* 2022; Zemanova 2020). Nonetheless, with the adoption of such a non-invasive method, it is imperative to use precise and efficient field methods to limit biases induced by misidentifications (Ashe & Hammond 2022; Johansson *et al.* 2020; Morrison *et al.* 2011).

In the computer vision domain, deep learning approaches have typically relied on very large training databases to achieve high recognition performance (LeCun *et al.* 2015; Liu *et al.* 2019; Wang & Deng 2021). However, the acquisition of large sample sizes in real-world scenarios is often quite challenging (Desai *et al.* 2022; Gómez-Vargas *et al.* 2023). This is particularly true in photo-identification studies, where the constraints are even more stringent. These studies inherently require individual-level information, yet often encounter few recapture occasions (Ashe & Hammond 2022; Meenakshisundaram *et al.* 2021; Suriyamongkol & Mali 2018). Moreover, numerous photos deemed of poor quality must be reviewed (Elliser *et al.* 2022; Urian *et al.* 2015). Generally, one picture is taken per capture (Bauwens *et al.* 2018; Salom-Oliver *et al.* 2022), or only the best one, if any, is retained (Elliser *et al.* 2022; Van Tienhoven *et al.* 2007). To address the issue of data scarcity, sophisticated protocols can be successfully designed to improve data collection at every encounter (Desai *et al.* 2022; Ferreira *et al.* 2020) or enhance the discrimination power of deep learning frameworks (Carter *et al.* 2014; Gómez-Vargas *et al.* 2023). While existing methods like Wild-ID and I3S Pattern demonstrate limitations in handling small training datasets, DL-ID maintains accuracy above 85%, even with as few as 3-4 picture samples per individual. This capacity holds great promise for its reliable implementation in the field, especially considering efforts to build a database that encompasses real-world challenges (Gómez-Vargas *et al.* 2023; Ravoor & T.s.b. 2020).

The introduction of a CI within the DL-ID framework corresponds to a recent approach to determine the likelihood that an image corresponds to a known or unknown individual, providing a quantitative measure of confidence in the identification process (Safaei *et al.* 2024). This additional safeguard is particularly essential in ecological and conservation studies, notably for monitoring programs whose final population or survival estimates can be subjected to large bias depending on the rate of misidentifications (Ashe & Hammond 2022; Johansson *et al.* 2020; Morrison *et al.* 2011). By assigning a CI, DL-ID introduces an extra element of transparency into the identification process, enabling researchers and conservationists to make more informed decisions relative to the match reliability and the study objective (Ashe & Hammond 2022; Elliser *et al.* 2022; Rigoudy *et al.* 2023). In our study, the ROC analysis reveals that DL-ID, with the incorporation of the CI, achieved higher AUC values compared to Wild-ID and I3S Pattern. This showcases the superior discriminative potential of DL-ID in distinguishing between true positive and false positive identifications and its robust discriminative power, even with small training dataset sizes.

The ensemble approach applied in DL-ID results in a substantial positive impact on the recognition performance (Malik *et al.* 2021; Mohammed & Kora 2023). In fact, by combining probabilities obtained from multiple instances of the model, DL-ID achieves an increase in individual identification accuracy from 53.2% to 63.1% on the most challenging training set (P_2_). This ensemble technique harnesses the collective strength of individual models, effectively reducing the risk of misidentifications and bolstering the model reliability in challenging scenarios. While our study employed a five-model ensemble for DL-ID, it is worth noting that resource availability can play a crucial role in determining the optimal ensemble size (Mohammed & Kora 2023). In challenging situations, such as small training dataset sizes (Gómez-Vargas *et al.* 2023) or individuals with subtle differentiation marks (Ferreira *et al.* 2020; Clapham *et al.* 2020), it may be beneficial to investigate the potential advantages of using larger ensembles. This could further enhance the model robustness and accuracy, particularly in scenarios where fine-grained distinctions are vital for accurate identifications (Malik *et al.* 2021). However, it is important to note that larger ensembles come with the drawback of potentially prohibitive training time and memory requirements, which should be carefully considered in resource-constrained environments (Mohammed & Kora 2023).

DL-ID also offers an advantage in terms of time efficiency, a critical factor for ecological research, particularly in large-scale and collaborative projects (Bauwens *et al.* 2018; Carpentier *et al.* 2016; Rigoudy *et al.* 2023; Schofield *et al.* 2019). This efficiency translates into accelerated image processing, enabling researchers to shift their focus towards the substantive aspects of data analysis and interpretation (Tuia *et al.* 2022). Compared to Wild-ID and I3S Pattern, DL-ID substantially reduces the time required for both training and inference phases. With a training duration of only two hours for the ensemble model and a processing time of only 10 seconds for 179 images during inference, DL-ID streamlines the workflow and enhances research productivity.

The adaptability of DL-ID is showcased not only in its application to Humboldt penguins but also in its seamless transfer to Hermann’s tortoises after training on a benchmark of similar size (P_80%_). The model’s intrinsic adaptability therefore extends beyond the boundaries of species similarity, a limitation often encountered in previous comparative studies (Matthé *et al.* 2017; Ravoor & T.s.b. 2020; Salom-Oliver *et al.* 2022). DL-ID’s versatility lies in its ability to be universally applied across various species and taxa with discernible and distinctive markings. Nevertheless, the dorsal shell of turtles was not investigated by previous photo-identification studies, which generally focused on face scutes (Carpentier *et al.* 2016; Carter *et al.* 2014) or ventral plastron (Salom-Oliver *et al.* 2022; Suriyamongkol & Mali 2018). The stability of these features over time remains largely unexplored outside of this framework (Bentley *et al.* 2021). While major changes are very unlikely due to the early calcification of the shell, its development under strong genetic control and physical properties at adulthood (Bentley *et al.* 2021; Zuffi *et al.* 2017), this requires a direct assessment to validate its suitability for long-term monitoring.

Accurate identification of individual animals throughout their life is vital for longitudinal studies (Bauwens *et al.* 2018; Schofield *et al.* 2019) and conservation efforts (Carpentier *et al.* 2016; Horn *et al.* 2014). The Hermann’s tortoise is already classified as a near threatened species by the UICN, and the Humboldt penguin is currently registered as vulnerable. Both species seem headed towards an endangered classification following improvements in their population monitoring and subsequent re-evaluation of their status (Graciá *et al.* 2020; Mcgill *et al.* 2022). In this context, implementing photo-identification methods that do not require capture is essential. Contrary to previous approaches relying on the turtle plastron (Salom-Oliver et al. 2022; Suriyamongkol & Mali 2018), DL-ID discriminates individuals based on their dorsal shell, eliminating the need for physical manipulations to perform identification. The current understanding of Humboldt penguins is limited, particularly in terms of accurate demographic trends and life history parameters (De la Puente *et al.* 2013; Mcgill *et al.* 2022). Additionally, there is an urgent need for detailed information on their dispersal and movements (Mcgill *et al.* 2022). Our new method for photo-identification using DL-ID provides a valuable tool that could contribute significantly to reliable monitoring and effective conservation of Humboldt penguins. Reliable long-term identification, even through molts, is crucial for in-depth spatial and population dynamics studies (Mcgill *et al.* 2022). Photo-identification with DL-ID offers a non-invasive, effective, reliable and cost-efficient solution, as previously shown in studies on the African penguin (Burghardt *et al.* 2004; Sherley *et al.* 2010). Additionally, zoological institutes (Burghardt *et al.* 2004; Desai *et al.* 2022; Núñez-López *et al.* 2021) and other captive settings (Carpentier *et al.* 2016; Ferreira *et al.* 2020) provide ideal conditions to collect sufficient ground truth data to train and test DL-ID models before their field application (Sherley *et al.* 2010).

Future research should include a broader spectrum of species, particularly those with diverse physical and behavioral characteristics, to better evaluate the adaptability of DL-ID. Refining the model’s capability to accurately identify individuals across time at various life stages is equally important. While it may not be suitable for certain species at specific life stages, enhancing its adaptability to investigate species that undergo considerable morphological transformations during their lifetime, such as amphibians and butterflies, would broaden the potential application of DL-ID. Indeed, improving this feature would significantly enhance its effectiveness for long-term studies and its utility in ecological conservation efforts. Therefore, subsequent research should concentrate on enhancing the adaptability and feature extraction abilities of the model. Key areas for improvement include boosting its accuracy and dependability by training on a diverse range of species and ecological environments. This is particularly important in settings characterized by significant variability in visual features and conditions. The development of tools for real-time identification and monitoring has the potential to transform conservation efforts, enabling faster and more effective responses in critical ecological scenarios. Such scenarios include tracking endangered species or managing invasive populations. Advancements in this area would not only open new perspectives of use, but also make a substantial contribution to the field of ecological conservation and research.

