## Appendices for "Deep learning-based photo-identification for non-invasive monitoring of animal populations: Application to penguins and tortoises"

### Appendix 1: DL-ID vs. popular DL methods

Currently, popular approaches to image identification include transfer learning and Siamese networks. In transfer learning, large convolutional networks pre-trained on state-of-the-art datasets like Imagenet are adapted for new tasks by replacing the final classifier layer to match the number of classes specific to the problem under study. The primary concept is to train this new classifier layer without altering the pre-trained weights of preceding layers. However, fine-tuning does allow certain earlier layers' parameters to be adjusted, potentially enhancing performance. In Siamese networks, there are three identical networks with shared parameters that are trained simultaneously. Instead of yielding a probability distribution for classes, the output is an embedding vector in a pre-defined dimensional space, where this linear dimension serves as a hyperparameter. The goal is to ensure that embedding images belonging to the same class are close according to a chosen metric in this space, another hyperparameter.

Both of these approaches have been effectively applied to image recognition projects. However, we believe that they do not offer significant advantages over our method for applications similar to those described in this work. In transfer learning, although deep layers can effectively capture general low level features from the original training set, most are not relevant for specialized tasks. The main advantage of this approach lies in leveraging a complex network trained on a large dataset like Imagenet, with approximately 14M images, to achieve high accuracy and low overfitting. Historically, the accuracy of convolutional architectures has improved as the networks became more complex, primarily due to advancements in parallel computing and GPU technology. Consequently, transfer learning remains a compromise in scenarios with limited computing resources.

Siamese networks are gaining attention for their performance in few-shot learning, particularly when labeled images for training are scarce. However, they present their own challenges. First, they are prone to overfitting when training data is sparse, which is one of the key focus areas of this article. Additionally, they require more computational resources due to their architectural complexity, which involves processing images through three network copies and minimizing a cost function over triangles in a vector space of a much higher dimension than used in the classifying top layer in a CNN. Another potential problem is that the usual cost functions used, based in the triplet loss metric, are not efficient during training and require precise tuning of a threshold and sometimes a semi-supervised selection of training examples using algorithms like online mining. Furthermore, adapting Siamese networks for label prediction requires creating class representative vectors in the embedding space and classifying new observations by choosing the closest representative, an operation that can be computationally intensive both during training and inference. Other inconveniences involve precise tuning of the dimension of the embedding space and the distance used for predicting classes since Siamese networks are sensitive to these two aspects.

In this work, we have instead chosen to use a pre-existing architecture that has performed well on Imaginet. Unlike transfer training, we trained this architecture from a complete random initialization, leveraging the current availability of consumer GPUs. Our approach achieved high accuracy results in both closed- and open-set recognition tasks. Comparisons between our method, transfer learning and Siamese networks using our datasets and the same basic network architecture show that the full training of a modern general purpose CNN works better in our usage case than these two alternatives, even with a limited number of images without augmentation.

**Appendix 2: I3S Pattern and Wild-ID analyzes**

Detailed technical information about the software programs is available both in their initial publication (Bolger *et al.* 2012; Van Tienhoven *et al.* 2007) and respective user manuals (Bolger *et al.* 2011; Den Hartog & Reijins 2014). It is specified that both software programs have been designed to help researchers in carrying out the recognition process and not to replace them, hence the limited capacities (Schneider *et al.* 2019; Schofield *et al.* 2019) and inherent subjectivity of human observers (Johansson *et al.* 2020; Urian *et al.* 2015) remain problematic. This section describes the main steps and specificities of the protocol used for the ***Known*** and ***New*** experiments on the Humboldt penguin. For both I3S Pattern and Wild-ID analyses, the same pictures were employed as in our DL-ID framework to allow a fair comparison of performance, except for a dedicated testing of I3S Pattern performed in optimal conditions (see below). Due to the functioning of both software programs, the training dataset was only acting as an identification database built from former photo shooting sessions (Matthé *et al.* 2017; Suriyamongkol & Mali 2018; Van Tienhoven *et al.* 2007), unlike deep learning approaches depending on and learning from this training material (LeCun *et al.* 2015; Schneider *et al.* 2019). To avoid transcription mistakes, gain significant time, and improve readability of the outcome, a Python script was written to compute the identification rate per individual and the total accuracy from their output files.

Concerning I3S Pattern analyzes, three reference points should be selected on each picture for rotation and scaling purposes (Den Hartog & Reijins 2014). The first one was positioned in the middle at the bottom of the black band, whereas the two others were placed at the junction between the chest and each foot (Appendix.3). For the ***Known*** experiment, two analyses were performed with slightly different framings (Appendix.2): 1. the optimal comparison - following the user guidelines, the mouse was used to crop only the spotted chest, leaving out all background and the black band ; 2. the direct comparison - in order to have results directly comparable with Wild-ID and our DL-ID framework, the entire picture was selected by placing a point at each corner. Owing to its higher accuracy, the optimal framing was used for the ***New*** experiment and time measurements. For every experiment, multiple tests comparing the dedicated training set to the validation set were run with the aim of identifying for each internal parameter the value, which optimizes the accuracy. As the most relevant variable (Den Hartog & Reijins 2014; Givord-Coupeau & Rey 2023), the number of key points extracted was first searched for an optimum and was established to be around 18 for the optimal comparison and 33 for the direct comparison. Then, following the parameter order on user interface, all others apart from color weights were successively set at five values, i.e. default value (dv), dv±25% and dv±50%, to quantify the impact on the identification. This means the parameter value maximising the accuracy was retained each time before pursuing the analysis further. In the end, only the best combination of parameters was used to perform the identification of pictures belonging to the testing set, and this procedure was repeated before every identification experiment (Appendices 5 and 5).

In the case of Wild-ID processing, the same cropped images as the other recognition methods were directly fed to the algorithm for key point extraction. Wild-ID is almost totally automated. Its user interface does not provide any access to enable modification of pictures and pre-defined internal parameters as well (Bolger *et al.* 2011), removing the opportunity to proceed with a validation step before the final test. Moreover, this software program cannot compare two datasets against each other, as it handles images in alphabetical order, and integrates each of them to the database inspected for matches after their own comparison. Therefore, a Python script was used to 1. provide the program with all pictures from the training dataset first and 2. integrate each testing image that was assigned with a random identifier in a second step. As the testing dataset remained unchanged across the ***Known*** experiment, pictures kept their identifier identical, avoiding introducing additional bias in analyses. As for the ***New*** experiment, pictures of new individuals were attributed a random identifier, so that all were treated after those of known penguins. Also, when a true match was proposed at top rank for a new penguin, the list was further inspected to determine the first non-matching image. Finally, based on the script created for I3S Pattern, a second code was written to fit the structure of Wild-ID output files, notably leaving out all lines concerning the training dataset.

| | nrElements | warpSize | relKey | maxAll | minRel | minRatio | accuracy |
|---|---|---|---|---|---|---|---|
| val_direct_P2 | 32 | 625 | 0.05 | 0.005 | 2.25 | 0.5 | **49.21** |
| val_direct_P5 | 28 | 500 | 0.05 | 0.007 | 1.5 | 0.5 | **57.59** |
| val_direct_P10 | 32 | 500 | 0.05 | 0.01 | 1.5 | 0.5 | **69.11** |
| val_direct_P20 | 32 | 500 | 0.05 | 0.01 | 1.5 | 0.5 | **74.87** |
| val_direct_P30 | 35 | 625 | 0.05 | 0.005 | 1.5 | 0.25 | **82.20** |
| val_direct_P40 | 30 | 750 | 0.05 | 0.005 | 1.5 | 0.25 | **83.25** |
| val_direct_P50 | 33 | 750 | 0.05 | 0.005 | 1.5 | 0.5 | **87.43** |
| val_direct_P60 | 33 | 750 | 0.05 | 0.005 | 2.25 | 0.5 | **86.91** |
| val_optimal_P2 | 17 | 500 | 0.063 | 0.007 | 1.875 | 0.375 | **46.60** |
| val_optimal_P5 | 20 | 500 | 0.025 | 0.013 | 1.5 | 0.5 | **83.77** |
| val_optimal_P10 | 19 | 500 | 0.025 | 0.01 | 1.5 | 0.375 | **89.01** |
| val_optimal_P20 | 18 | 500 | 0.037 | 0.01 | 1.875 | 0.5 | **94.24** |
| val_optimal_P30 | 18 | 500 | 0.037 | 0.01 | 1.5 | 0.5 | **95.81** |
| val_optimal_P40 | 18 | 500 | 0.037 | 0.01 | 1.5 | 0.5 | **95.81** |
| val_optimal_P50 | 18 | 500 | 0.037 | 0.01 | 1.5 | 0.25 | **96.86** |
| val_optimal_P60 | 18 | 500 | 0.037 | 0.01 | 1.875 | 0.25 | **97.38** |

**Appendix 3:** I3S Pattern validation step for Known experiment. For both comparison methods and all training sizes, the final combinaison of settings and the identification rate are displayed.

| | nrElements | warpSize | relKey | maxAll | minRel | minRatio | AUC score |
|---|---|---|---|---|---|---|---|
| val_optimal | 15 | 500 | 0.05 | 0.01 | 1.5 | 0.5 | 0.928 |
| val_optimal | 18 | 500 | 0.05 | 0.01 | 1.5 | 0.5 | 0.927 |
| val_optimal | 20 | 500 | 0.05 | 0.01 | 1.5 | 0.5 | 0.912 |
| val_optimal | 18 | 500 | 0.037 | 0.01 | 1.5 | 0.5 | 0.934 |
| val_optimal | 18 | 500 | 0.037 | 0.01 | 1.875 | 0.5 | 0.938 |
| val_optimal | 18 | 500 | 0.037 | 0.01 | 1.875 | 0.25 | 0.947 |
| **test_optimal** | **18** | **500** | **0.037** | **0.01** | **1.875** | **0.25** | **0.931** |
| val_direct | 31 | 500 | 0.05 | 0.01 | 1.5 | 0.5 | 0.833 |
| val_direct | 33 | 500 | 0.05 | 0.01 | 1.5 | 0.5 | 0.837 |
| val_direct | 35 | 500 | 0.05 | 0.01 | 1.5 | 0.5 | 0.84 |
| val_direct | 33 | 750 | 0.05 | 0.01 | 1.5 | 0.5 | 0.87 |
| val_direct | 33 | 750 | 0.05 | 0.005 | 1.5 | 0.5 | 0.875 |
| val_direct | 33 | 750 | 0.05 | 0.005 | 2.25 | 0.5 | 0.887 |
| **test_direct** | **33** | **750** | **0.05** | **0.005** | **2.25** | **0.5** | **0.836** |

**Appendix 4:** I3S Pattern validation step for New experiment. For both comparison methods, settings are explored using the order on I3S Pattern user interface to optimize the AUC score. Validation steps are performed on the P60% set and final test is using the P80% dataset.

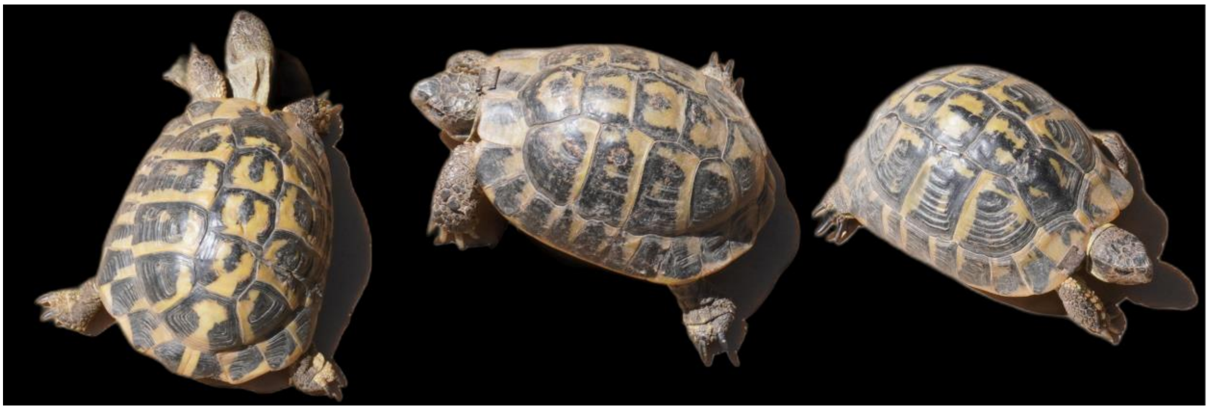

**Appendix 5:** Image sample of three Hermann's tortoises, corresponding to the format provided to DL-ID for processing. Pictures are rescaled and most of their background is automatically removed, leaving only small shadow parts.

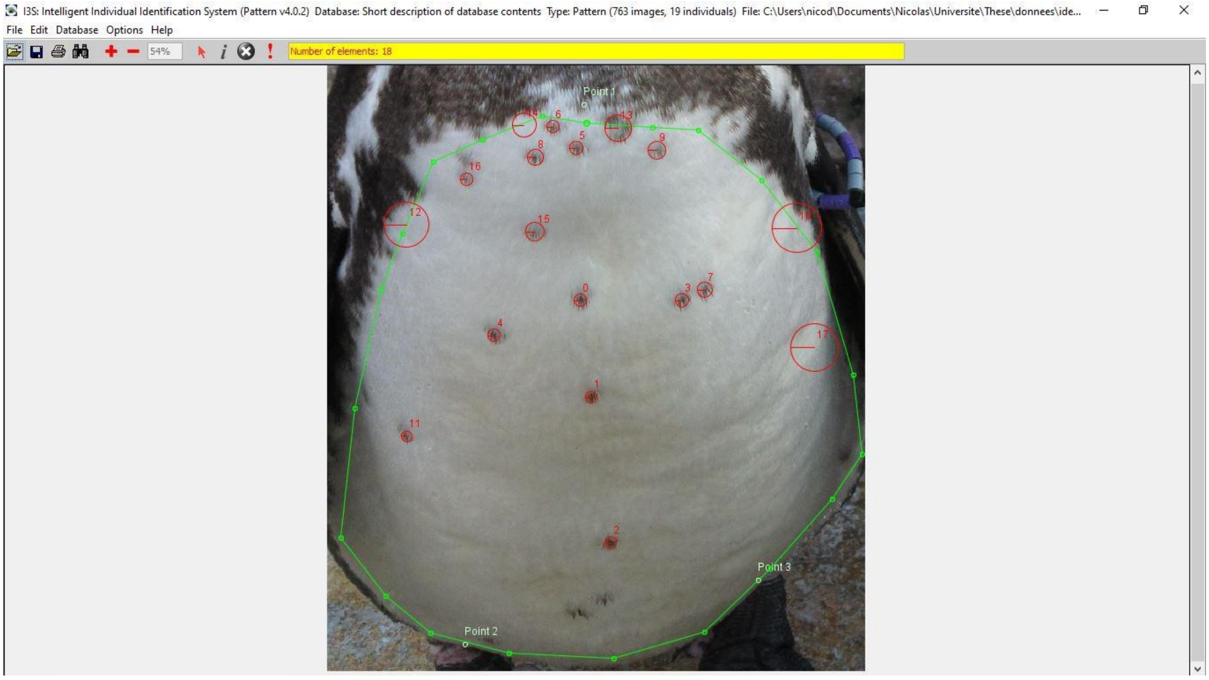


**Appendix 6:** Overview of I3S Pattern interface, displaying the 3 points used as references for scaling, the region of interest and the 18 points extracted from there for identification purpose.

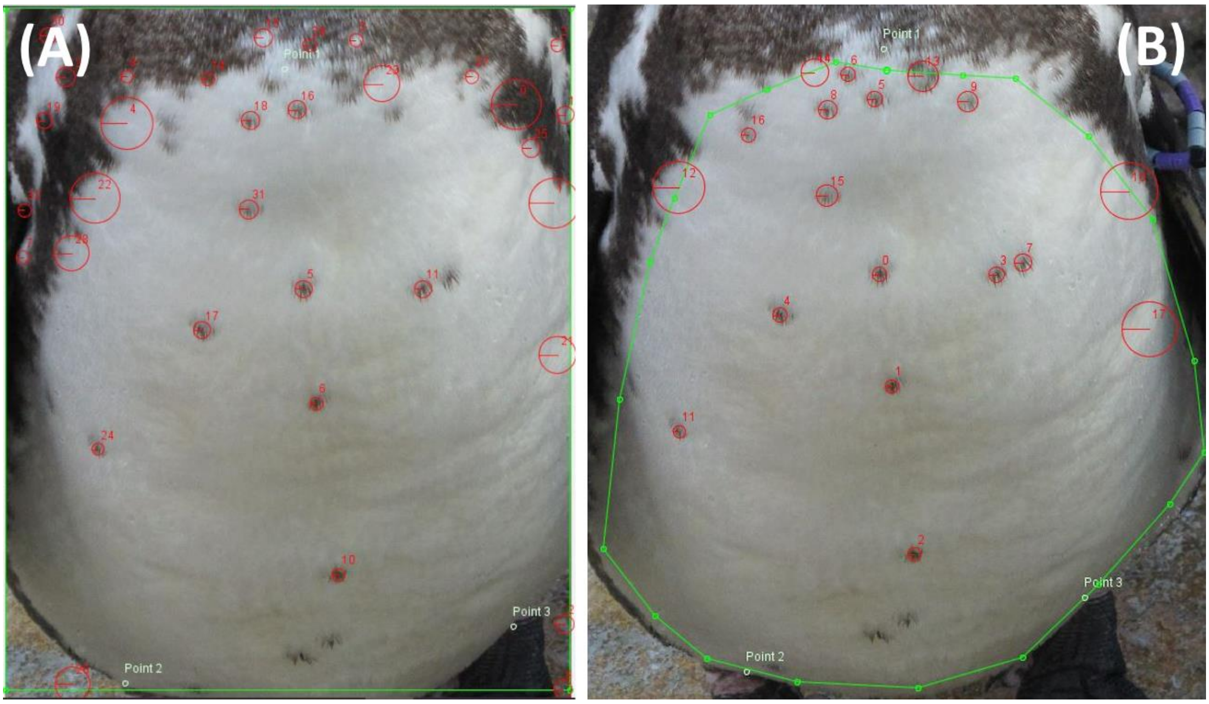


**Appendix 7:** Identification experiments performed on I3S Pattern, using different framing and optimized number of extracted points. For the direct comparison, the full picture is selected to obtain performance comparable with other methods (A). For the optimal comparison, the program is used as intended by its developers, with careful cropping around the chest and its spot pattern (B).
